# Photoselective sequencing: microscopically-guided genomic measurements with subcellular resolution

**DOI:** 10.1101/2022.09.12.507600

**Authors:** Sarah Mangiameli, Haiqi Chen, Andrew S. Earl, Julie Dobkin, Daniel Lesman, Jason Buenrostro, Fei Chen

## Abstract

In biological systems, spatial organization is interconnected with genome function and regulation. However, methods that couple high-throughput genomic and epigenomic profiling with spatial information are lacking. Here, we developed Photoselective Sequencing, a spatially-informed DNA sequencing method to assay collections of cells or subcellular regions that share a unifying morphological trait. In Photoselective Sequencing, we prepare a blocked fragment library within a fixed biological specimen. Guided by fluorescence imaging, we remove the block in specific regions of interest using targeted illumination with near-UV light, ultimately allowing high-throughput sequencing of the selected fragments. To validate Photoselective Sequencing, we profile chromatin openness in fluorescently-labeled cell types within the mouse brain and demonstrate strong agreement with published single-cell ATAC-seq data. Using Photoselective Sequencing, we characterize the accessibility profiles of oligodendrocyte-lineage cells within the cortex and corpus-callosum regions of the brain. We develop a new computational strategy for decomposing bulk accessibility profiles by individual cell types, and report a relative enrichment of oligodendrocyte-progenitor-like cells in the cortex. Finally, we leverage Photoselective Sequencing for unbiased profiling of DNA at the nuclear periphery, a key chromatin organizing region. We compare and contrast the Photoselective Sequencing profile with lamin ChIP-seq data, and identify features beyond lamin interaction that are correlated with positioning at the nuclear periphery. These results collectively demonstrate that Photoselective Sequencing is a flexible and generalizable platform for exploring the interplay of spatial structures with genomic and epigenomic properties.

## Main

The genome encodes the information that underlies cell state and function in its sequence and structure. This structure is organized across diverse length scales. At the subcellular scale, the arrangement of the genome and associated proteins within the volume of the nucleus is a central regulator of gene expression programs^1–4^. This includes transcriptional activation by enhancer-promoter interactions or sequestration of biologically inactive sequences to heterochromatic domains throughout the nucleus^5,6^. Within complex tissues, cells respond to environmental cues or intercellular signaling through diverse transcriptomic and epigenomic states^7–9^. A complete understanding of the interplay between spatial structures and sequence-based information will require new methods that simultaneously measure these properties.

An emerging strategy is to perform a genomic or epigenomic measurement within particular spatial regions under the guidance of microscopic visualization^10–13^. For example, laser-capture microdissection and patch pipette aspiration are early implementations of this strategy where the relevant subpopulation of cells is physically excised from the sample prior to downstream sequencing-based analysis^14–16^. However, these methods do not have subcellular spatial resolution, are subject to contamination from neighboring cells, and can be laborious if large numbers of individual cells are needed. A similar class of methods use photoactivation to capture nucleic acids from specific nuclei within a sample^10^. However, the methods largely apply to transcriptomic measurements, are only suitable for analyzing entire nuclei (i.e not mitochondrial or other extranuclear DNA), and do not provide sub-cellular resolution.

To address these shortcomings, we introduce Photoselective Sequencing (PSS), a new method for sequencing DNA from spatial regions of interest (ROI) within a fixed biological specimen (Fig. 1A). PSS breaks down into four basic stages: (1) construction of a photocaged DNA-fragment library *in situ*, (2) selective uncaging of the library using targeted illumination with near-UV light, (3) sample digestion and library purification, and (4) amplification of uncaged fragments followed by readout on an Illumina platform (Fig. 1A). During the *in situ* library construction stage, we use Tn5 transposase to produce a library of DNA fragments that are flanked by adapter sequences (tagmentation). Direct transposition of the sample assays chromatin openness (ATAC-seq library; Fig 1A, Step 1)^17^. However, transposase-mediated library preparations have been adapted for both unbiased and protein-targeted fragmentation *in situ*^*18,19*^.

**Fig. 1:**
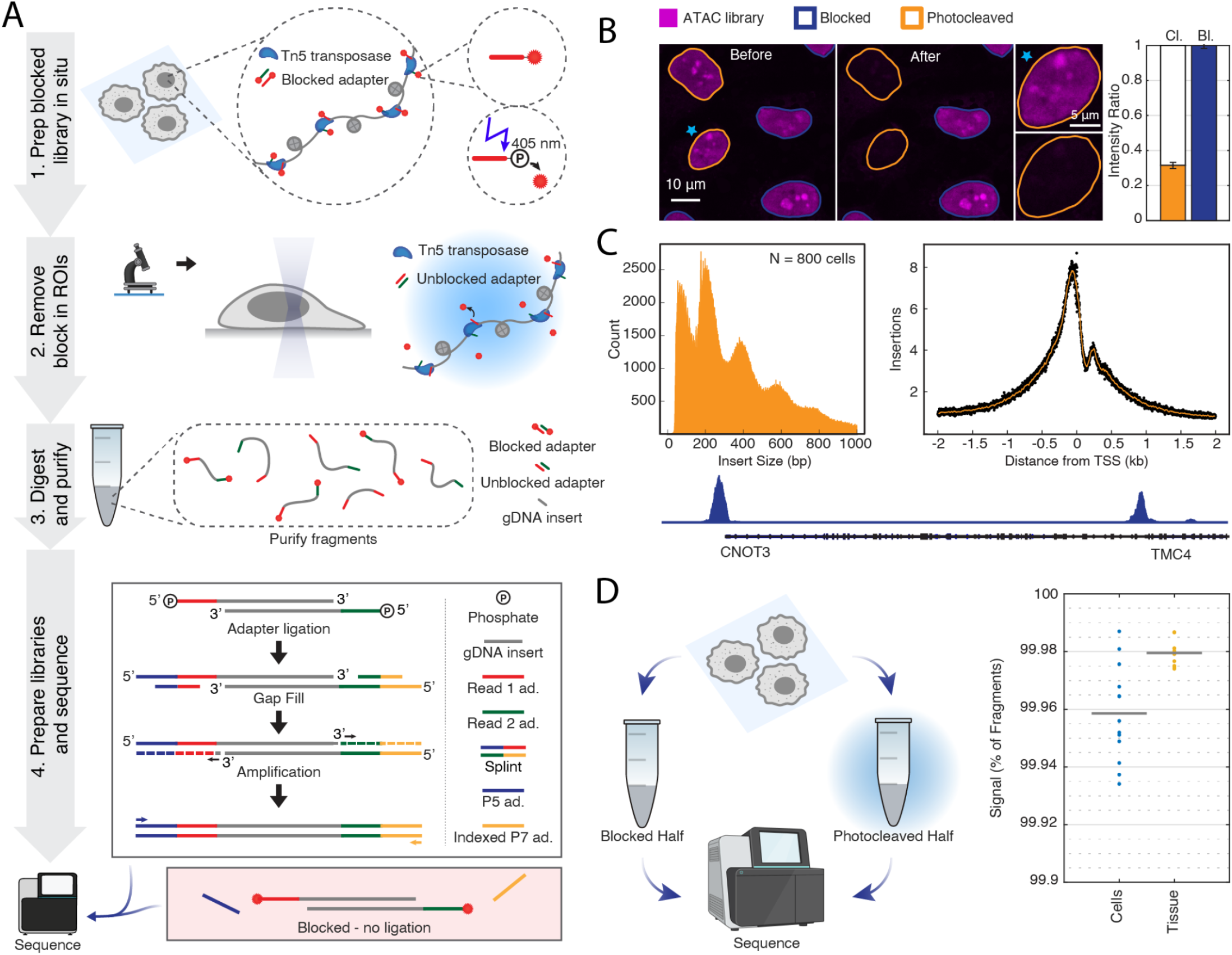
PSS enables genomic and epigenomic profiling with high spatial resolution. A) Schematic diagram showing the four-stage PSS protocol. In the first stage, we prepare a blocked library within an in-tact biological specimen that was previously stained for any markers necessary to recognize ROIs. To do this, we treat the sample with Tn5 Transposase (blue) that carries photocleavable blocked adapters (red and green). Tn5 transposase simultaneously fragments the genome and ligates the blocked PSS adapters to the ends of each segment (middle circle). The PSS adapter (top right circle) consists of an oligonucleotide sequence (red line), a photocleavable spacer (black line), and a fluorophore for visualization (red star). Exposure of the PSS adapter to near-UV light breaks the photocleavable linker, removing the fluorophore and exposing a 5’ phosphate group that enables subsequent steps of the library preparation. In the second stage of PSS, we visualize the specimen on the microscope and programmatically or manually identify ROIs. ROIs are specifically illuminated with near UV-light, removing the fluorophore and uncaging the library within these spatial regions. In the third stage, we digest the sample and purify the fragment libraries, resulting in a soluablized mixture of caged and uncaged fragments. Finally, we ligate secondary adapters to the 5’ phosphates of uncaged fragments (yellow and blue lines). The red panel illustrates a blocked fragment which does not have exposed 5’ phosphate groups and is unable to pass the ligation stage. A gap-fill step produces complete double stranded fragments. We subsequently amplify the library from these ligated adapters prior to next-generation sequencing. B) Uncaging ATAC libraries using targeted illumination, demonstrated in HeLa cells. The magenta color represents the fluorescence intensity of the fragment library, visualized by the 5’ fluorophore carried by the caged PSS adapters. Targeted illumination of the yellow-outlined nuclei releases the fluorophore from the adapter (after image) and results in a ∼70% decrease in fluorescence intensity (bar graph), indicating successful uncaging of the library. Non-targeted cells (blue outlines) do not decrease in intensity (bar graph), indicating that PSS is spatially precise. The starred cell is shown in the zoomed-in image to the right. C) Quality metrics for ATAC-seq libraries generated using the PSS method. Libraries show a characteristic periodic insert size distribution (histogram, left) due to histone positioning and a high insertion frequency into transcription start sites (right). Coverage traces show strong peaks upstream of highly-expressed gene loci (bottom trace). D) Left: schematic diagram illustrates the experimental process for the signal-to-noise calculation. Right: plot shows the percent of fragments that are counted as signals for cells and tissues. Dots represent the signal-to-noise ratio for biological replicates. Solid line indicates the mean.

To visualize the fragment library and reversibly block amplification, we use customized tagmentation adapters conjugated to a fluorophore using a photocleavable spacer (Fig. 1A, step 1). We next visualize the sample by microscopy and use fluorescent stains to guide ROI identification (Fig. 1A, step 2). Automated in-line image segmentation enables us to scalably assay thousands of individual ROIs localized throughout the sample. Selective illumination of the ROIs using a 405 nm laser line cleaves the fluorophores from the blocked adapters, revealing a 5’ phosphate group. After purification of the library from the sample (Fig. 1A, step 3), indexed secondary adapters containing priming sites for library amplification are ligated only to fragments where the 5’ phosphate groups were previously exposed through photocleavage (Fig. 1A, step 4). Caged fragments do not have available 5’ phosphates and cannot pass the ligation stage (Fig 1A, step 4, red box). Successfully-ligated fragments undergo a gap fill step and are amplified from the secondary adapters to produce sequencing-competent fragments which are read out an Illumina sequencer (Extended Data Fig. 1A).

## Results

### Validating the PSS workflow

To validate the PSS workflow, we first characterized ATAC-seq libraries in adherent cultured cells. To do this, we treated fixed and permeabilized HeLa cells with freely diffusing Tn5 transposase, which preferentially inserts into open chromatin regions^17,20,21^. Library generation was verified by visualization of the fluorescent label on the blocked tagmentation adapters (Fig. 1B, left image). When a subset of nuclei were exposed to near-UV light by scanning a focused 405 nm laser through these regions (Fig. 1B; Extended Data Fig. 1A-B and Movie 1), we observed an accompanying 80% decrease in fluorescence intensity within the exposed nuclei (Fig. 1B, bar graph). We next examined the sequencing characteristics of ATAC-seq libraries generated through the PSS workflow. To do this, we uncaged all fragments within the nuclei of 800 individual HeLa cells before digesting the sample and purifying the libraries. We ligated secondary adapters to the unblocked fragments via the free 5’ phosphate group, and subsequently used these adapters as PCR handles to generate the final sequencing libraries. After aligning the sequencing reads to the human genome, we observed a periodic fragment size distribution, a hallmark property of ATAC libraries that indicates sensitivity to nucleosome positioning (Fig. 1C, left). Furthermore, we observed a high insertion frequency across transcription start sites (TSS) genome wide, as indicated by a strong peak in the TSS enrichment plot (Fig 1C, right; Methods).

To assess the efficacy of the PSS blocking mechanism, we characterized the expected frequency of spurious fragments (see Methods - Signal to noise ratio calculation). Briefly, we constructed and purified PSS libraries using wells of HeLa cells or mouse brain slices. Once in solution, the libraries were split into paired blocked and photocleaved fractions and sequenced on an Illumina platform (Fig. 1D, schematic). We took the expected number of spurious fragments in the photocleaved library as the number of sequencing-competent fragments discovered in the blocked library. By this metric, we found that approximately 99.95% and 99.85% of detected fragments are signal in cells and tissues, respectively (Fig. 1D, plot), demonstrating the sensitivity of the PSS method.

### PSS reproduces aggregate single-cell ATAC-seq data in the mouse brain

To validate and demonstrate the PSS method, we generated chromatin accessibility profiles for both the dentate gyrus region and oligodendrocyte-lineage cells within the mouse brain (Fig 2A). The dentate gyrus region can be identified through tissue morphology and is composed primarily of hippocampal granule cells (Fig 2B, top image). Conversely, oligodendrocyte-lineage cells, which include mature oligodendrocytes and oligodendrocyte precursor cells, are scattered throughout the entire brain, and are identifiable via an immunostain against OLIG2 (Fig. 2B, bottom image). We selected these targets to demonstrate photoselection on various morphological traits, and because single-cell ATAC-seq (scATAC-seq) data are readily available for benchmarking the accessibility profiles of the associated cell types^22,23^.

**Fig. 2:**
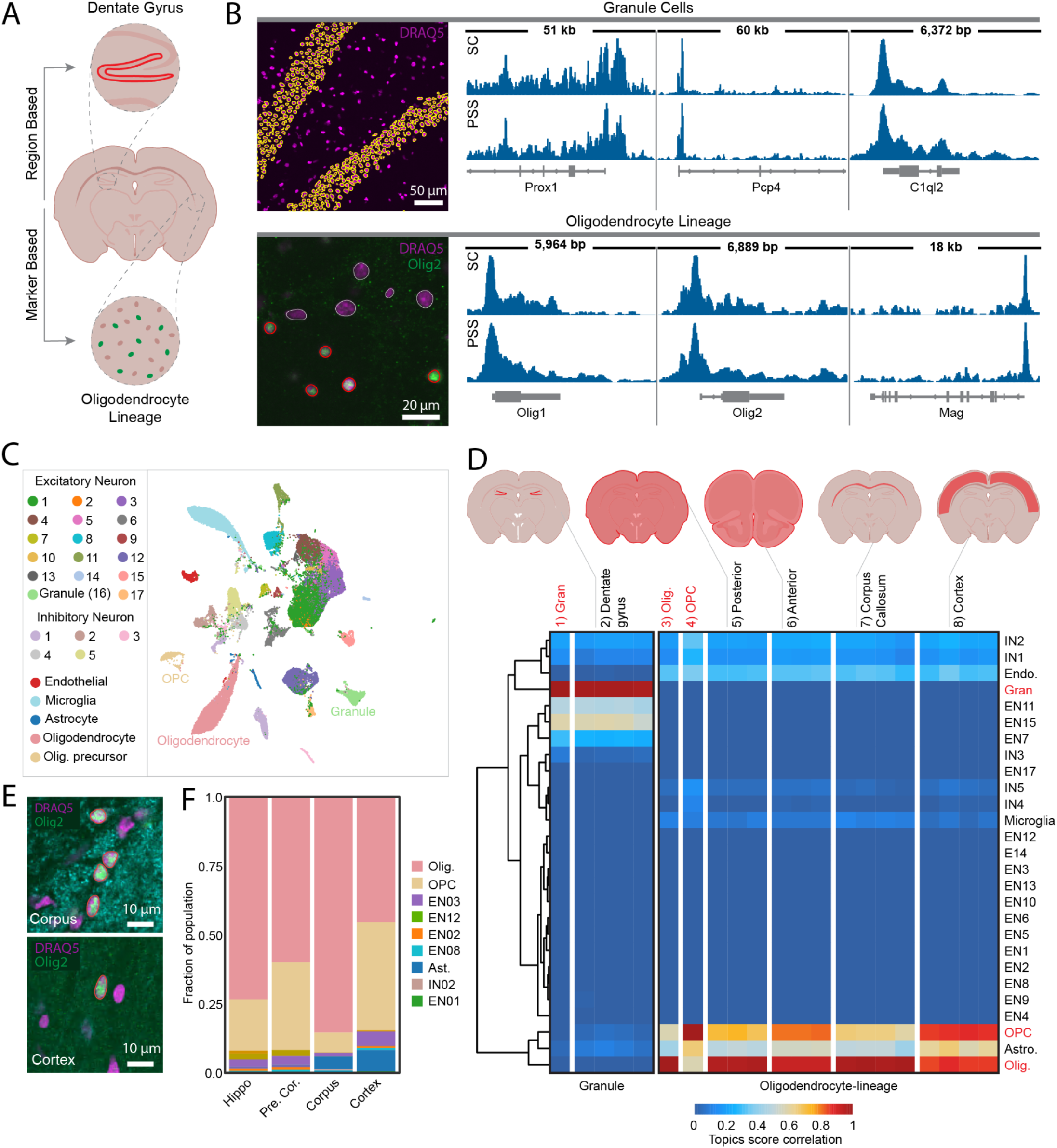
PSS robustly reproduces aggregate single-cell ATAC data and characterizes the regional composition of oligodendrocyte-lineage cells in the mouse brain. A) Morphologically guided selection of specific cell types in the mouse brain. The dentate-gyrus granule cells are resident to a characteristic v-shape region in the mouse hippocampus (top). OLIG2 immunofluorescence marks oligodendrocyte-lineage cells (bottom). B) Example images show target ROIs for the dentate gyrus, identified using a DRAQ5 DNA stain (purple), and oligodendrocyte-lineage cells, identified OLIG2 immunofluorescence (green). Yellow (granule cells) or red (oligodendrocyte-lineage cells) outlines indicate targeted cells. Coverage traces (right) compare the PSS ATAC-seq profiles (bottom traces; PSS) to aggregated single-cell ATAC-seq profiles for the corresponding cell types (top traces; SC) at example marker gene loci. C) UMAP plot showing clustering of reference scATAC-seq data on cisTopic z-score. D) Heatmap shows pairwise Pearson correlation coefficients of the cisTopic z-scores for the PSS data and aggregate scATAC-seq profiles. Red text indicates pseudobulk scATAC-seq data corresponding to a PSS-targeted cell type. Schematic diagram shows brain region for PSS samples. E) Photoselective sequencing compares oligodendrocyte-lineage cells in the cortex and corpus callosum. Schematic inserts indicate the corpus callosum and cortex regions of the hippocampus, while images show example target oligodendrocyte-lineage nuclei within the specified region. F) Cell type decomposition for oligodendrocyte-lineage PSS samples shows that white-mater rich tissues contain more mature oligodendrocytes.

To implement PSS for oligodendrocyte-lineage and granule cells, we collected a series of 10 micron tissue sections from a fresh-frozen mouse brain (Extended Data Fig. 2A; Methods). For dentate gyrus selection, sections were acquired from the hippocampal region. To investigate any region-specific differences in oligodendrocyte-lineage cells, sections were collected from both the anterior and posterior brain regions. Tissue sections for oligodendrocyte-lineage cell selection were immunostained for OLIG2, and all sections underwent the *in situ* phase of the PSS library preparation (Fig. 1A, step 1), and were stained with DRAQ5 (far-red DNA stain) to visualize nuclei. During the photoselection phase (Fig. 1A, step 2), nuclei within the dentate gyrus region were automatically detected based on proximity to neighboring nuclei (n=4 biological replicates; Fig. 2B, yellow outlines; Methods - Photoselection). Similarly, oligodendrocyte-lineage cells were detected algorithmically by identifying nuclei with a high fluorescence intensity from the OLIG2 immunofluorescent marker (n=6 biological replicates; Fig. 2B, red outlines; Methods - Photoselection). We selectively illuminated the target nuclei with near-UV light and then completed the remainder of the library preparation (Fig. 1A, steps 3-4), including readout on an Illumina platform.

For validation, we compared the PSS accessibility profiles to those of aggregated granule cells and oligodendrocytes from an annotated single-cell ATAC-seq (scATAC-seq) data set^22^. Visual comparison of the scATAC-seq and PSS profiles at marker gene loci for the respective cell types suggest a strong agreement between the two approaches (Fig. 2B, traces; Methods - Smoothed coverage traces). To quantitatively characterize the correlation between the PSS and scATAC-seq data genome wide, we first performed cisTopic analysis on the scATAC-seq data to generate a reference set of cis-regulatory control regions (topics) for further analysis (Methods - cisTopic analysis)^23,24^. We show the clustered scATAC-seq data (based on cisTopic z-score) in the UMAP representation, with the relevant clusters annotated (Fig. 2C). We next leveraged the topics regions to jointly analyze the PSS and scATAC-seq data. To do this, we quantified the cisTopic z-scores for each PSS replicate and the cluster-aggregated scATAC-seq data, calculated the pairwise Pearson correlation coefficients, and visualized the results as a hierarchically-clustered heat map (Fig. 2D, sections 1-6). The results show a high level of reproducibility between replicate PSS samples, and that the PSS accessibility profiles strongly correlate with those of the corresponding cell types in the single-cell data (see Fig. 2D, sections 1,3 and 4 for single-cell reference data).

Notably, the oligodendrocyte-lineage cell accessibility profiles from anterior (∼Bregma 2.2mm, Cortex and Anterior Olfactory Nucleus) and posterior (∼Bregma ∼1.6mm, Hippocampal Formation, Cortex, Thalamus, and Hypothalamus) contained signatures of both mature oligodendrocytes and oligodendrocyte precursor cells, but in different proportions (Fig. 2D, compare sections 3-6). This result suggests a relative enrichment of mature oligodendrocytes in the posterior section compared to the anterior section. We hypothesized that this difference in cell-type composition reflects the functional properties of the associated brain regions. In particular, mature oligodendrocytes are responsible for producing myelin, a lipid based material that sheaths and insulates nerve axons throughout the central nervous system^25,26^. In particular, we observed increased white matter regions (e.g. corpus callosum) and other fiber tracts in the posterior section, and we thus sought to investigate if mature oligodendrocytes comprise a higher proportion of oligodendrocyte-lineage cells in white-matter tracts vs cortical regions.

### PSS compares oligodendrocyte-lineage cell composition in the cortex and corpus callosum regions of the mouse brain

To test if the observed differences in oligodendrocyte-lineage cell composition between anterior and posterior tissues sections could be explained by tissue composition, we applied PSS to examine oligodendrocyte-lineage cells within white and gray matter subregions of a coronal brain section. Specifically, we generated ATAC-seq profiles for oligodendrocyte-lineage cells within the corpus callosum, a connective white matter region consisting millions of myelinated axons, and the cortex, the outermost gray matter layer of the brain (Fig. 2E, Extended Data Fig 2B). To do this, we collected tissue sections (n=8 biological replicates) from the mouse hippocampus, which were stained for OLIG2 and processed according to the in *situ library* generation phase of the PSS protocol (Fig 1A, step 1). During the imaging stage, we acquired whole-section scans to visualize the overall tissue morphology and identify the cortex and corpus callosum regions. We targeted either oligodendrocyte-lineage cells in either the cortex (n=4 biological replicates) or corpus callosum (n=4 biological replicates) during the selective illumination phase (Fig 1A, step 2), and proceeded through sequencing.

We compared the corpus-callosum and cortex oligodendrocyte-lineage cell profiles to the scATAC-seq data by calculating the cisTopic z-score correlations as described above (Fig. 2D, section 7-8). As expected, we found that both brain regions contain signatures of mature oligodendrocytes and oligodendrocyte precursor cells. However, oligodendrocyte-lineage cells from the corpus callosum region correlated more strongly with mature oligodendrocytes, consistent with our previous observation. To expand on this result, we leveraged the scATAC-seq data to decompose the PSS oligodendrocyte-lineage cell accessibility profiles by cell type. To do this, we performed stepwise linear regression on the cisTopic z-scores with the pseudobulk scATAC-seq profiles as independent variables (Methods - Stepwise linear regression). The regression coefficients were taken as the proportions of the constituent cell types in the PSS oligodendrocyte-lineage cell data. We found that mature oligodendrocytes comprise 73.4% of the oligodendrocyte-lineage cell population posterior tissue sections compared to 60.0% in anterior tissue sections. This effect intensifies in pure gray and white matter regions, where we estimate 85.5% of corpus-callosum oligodendrocyte-lineage cells are mature oligodendrocytes, compared with 45.5% in the cortex (Fig. 2F). These results demonstrate that PSS sensitively detects distinct epigenomic profiles within specific regions of the mouse brain. Furthermore, our method for co-analyzing the PSS data with existing scATAC-seq atlases enables decomposition of the bulk PSS accessibility profile into the contributions of individual cell types. Using this combined experimental and computational framework we demonstrate that PSS can infer the spatial distribution of specific cell types across tissue regions.

### PSS assays genomic sequences associated with the nuclear periphery

We next sought to define sub-cellular structures using PSS. The nuclear periphery is a key structural component of the nucleus known to play an important role in genome organization and gene regulation^27–29^. Furthermore, misregulation of interactions between the genome and the nuclear periphery have been implicated in age-related disorders and disease^28^. Therefore, a comprehensive understanding of the features that influence genome-nuclear periphery interactions would elucidate basic biological principles and have relevance in the context of human wellbeing. Previous studies have investigated specific protein-DNA interactions at the nuclear periphery, and in particular identified hundreds of lamin associated domains that range from 0.1-10 Mb in size^30^. Typically, these domains are defined by profiling Lamin B1-DNA interactions, either by proximity labeling assays or by chromatin immunoprecipitation followed by high throughput sequencing (ChIP-seq)^31–34^. However, these assays measure specific protein-DNA interactions and do not probe all sequences at the nuclear periphery in an unbiased manner. Using PSS, we selectively sequence the DNA at the nuclear periphery. We compare our results to published lamin B1 ChIP-seq data^35^ while exploring general features of peripheral DNA.

We constructed PSS libraries in adherent human fibroblast cells (IMR90) cells that were previously immunostained against Lamin B1 as a marker for the nuclear periphery (Fig. 3A, green), with an important modification. Specifically, we performed a gentle histone disruption treatment to facilitate unbiased Tn5 transposase insertion across all chromatin states (Methods - Unbiased Tn5 transposase insertion)^36^. During the photoselection phase, we unblocked DNA fragments at the nuclear periphery (n=3 biological replicates, 4000 cells each), defined as the two-dimensional boundary of the nucleus in the focal plane where the nuclear area is maximized (Fig 3A, yellow outlines; Methods - Photoselection). Once purified, the libraries from each sample were split into paired halves, one of which was prepared as an input library to enable enrichment-based measurements during the analysis stage (Methods - Enrichment Traces).

**Fig. 3:**
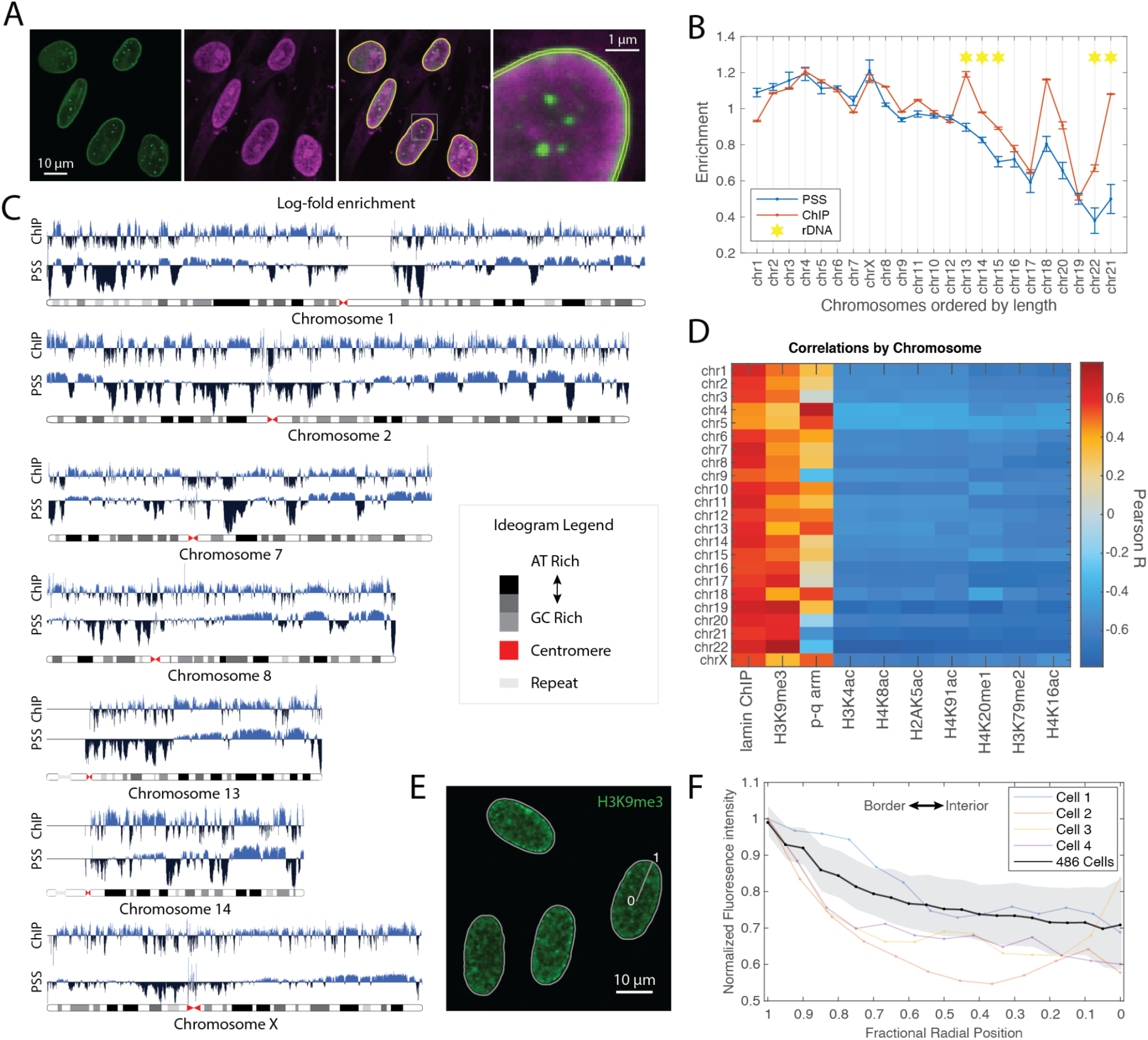
Photoselective sequencing identifies features of chromatin at the nuclear periphery. A) Images show lamin immunostain (green), an unbiased PSS library (magenta), and the merged image. Yellow outlines indicate the nuclear periphery region. Enlarged image illustrates the 300 nm -thick target region. B) PSS nuclear periphery profile compared to a published LamB ChIP at the chromosome level (chromosomes ordered by length). Plot shows fold-enrichment for PSS or ChIP-seq compared to an input (untargeted) library. Error bars represent the standard deviation (2 ChIP replicates, 3 PSS replicates). Starred chromosomes contain rDNA repeats. C) Log-fold enrichment profiles for nuclear periphery PSS and LamB ChIP-seq for a subset of chromosomes (see Extended Data Fig. 3 for all chromosomes). Profiles are pooled across replicates. D. Heatmap correlates PSS profiles to ChIP-seq profiles of histone modifications from the ENCODE database on a per chromosome basis. Chrs 4-5 profiles are particularly correlated with distance from the centromere (Extended Data Fig. 3), and all chromosomes show a positive correlation with the H3K9me3 ChIP-seq profile. E) Immunostain against H3K9me3 shows enrichment at the nuclear periphery in example cells. F) Radial fluorescence intensity profile for 468 cells immunostained for H3K9me3. Black line indicates the mean fluorescence intensity while the shaded gray area represents the standard deviation. Colored traces show the radial intensity profiles for the four example cells in E).

### Nuclear Periphery PSS enrichment is inversely correlated with chromosome size

We first asked which chromosomes are most enriched for reads in the PSS and ChIP-seq data (Methods - Enrichment traces). We found that the enrichments are generally inversely correlated with chromosome length, consistent with previous observations that the larger chromosomes are more peripheral on average (Fig. 3B)^37,38^. However, there is a clear discrepancy between the PSS and ChIP-seq enrichments for chromosomes 13-15,19, 21 and 22. Notably, with the exception of chromosome 19, these chromosomes each harbor nucleolar organizing regions, and thus interact with the nucleoli (Fig. 1B, starred). Besides having a key structural role in the nuclear lamina, lamin B1 protein is present throughout the nucleoplasm (Fig 1A., green) and contributes to nucleolar organization^39^. The relatively higher ChIP-seq enrichment for these chromosomes may stem from interactions with nucleoplasmic and nucleolar lamin B1 protein, which do not occur at the periphery and are not detected by PSS. To remove the effects of chromosome positioning, the lamin B1 ChIP-seq and PSS profiles are normalized on a per chromosome basis for all further computation.

### PSS identifies characteristic features of sequences that localize to the nuclear periphery

We next compared the lamin B1 ChIP-seq and PSS profiles genome wide. To do this, we calculate the log-fold enrichment along the coordinates of the genome using a 100 kb bin size (Fig. 3C; Extended Data Figure 3; Methods - Enrichment traces). We selected this bin size because it’s on the scale of the smallest lamin associated domains, and because replicate PSS profiles were strongly correlated using this binning (Extended Data Fig. 4A). We observe that the PSS and ChIP-seq profiles display many shared features and are strongly correlated (r=0.62 on average) genome wide (Fig. 3C and D). Notable exceptions include chromosomes 4 and 5, which show a relatively low correlation (r=0.44 and 0.45 respectively). Upon further investigation, we found that the PSS profiles for these chromosomes are strongly correlated with positioning along the chromosome, with depletion in the p-arm and enrichment in the q-arm (Fig. 3D; Extended Data Figure 3). One potential explanation for this effect is that these chromosomes are polarized such that the p-arms are oriented towards the nuclear interior on average, and thus less likely to interact with the periphery. Alternatively, the p-arms of these chromosomes may interact with the 3D surface of the nuclear lamina, but not the 2D border.

We next asked which histone modifications are characteristic of peripheral chromatin. Using the ENCODE database^40^, we analyzed all available histone modification ChIP-seq profiles that were collected from IMR90 cells. Binned enrichments were calculated for the ENCODE data described above, and correlated with the PSS data (Fig. 3D; Methods - Correlation analysis). We find that the PSS profile is positively correlated with the repressive mark H3K9me3 (r=0.47), and negatively correlated with active marks. To understand whether H3K9me3 modifies chromatin at the nuclear periphery that does not directly interact with lamin B1, we defined residuals by regressing the lamin B1 ChIP-seq profile out of the PSS profile (Methods - Correlation Analysis). We found that the H3K9me3 repressive mark was still correlated with the PSS residuals, even after removing the correlation with lamin B1 (Extended Data Fig. 4B). To validate our findings, we visualized the spatial localization of H3K9me3 using immunofluorescence (Fig 3E). We calculated the radial intensity profile of the fluorescent marker, and found a relative enrichment at the nuclear periphery as expected (Fig 3F; Methods - H3K9me3 radial intensity profile). Overall, our results suggest that H3K9me3-marked chromatin preferentially localizes to the nuclear periphery, even without direct lamin B interaction. Moreover, these results demonstrate PSS is able to probe subcellular nuclear organization, and sample orthogonal and complementary spatial relationships as compared to ChIP-seq, which samples molecular interactions.

## Discussion

PSS is a new method that assimilates imaging and sequencing data by enabling genomic or epigenetic sequencing measurements within specific ROIs, as visualized by fluorescence microscopy. We demonstrate that PSS flexibly applies across ROIs that span order-of-magnitude size scales, from entire regions of the mouse brain to the nuclear periphery in IMR90 cells. In-line image segmentation enables ROI generation based on an extensive range of spatial features and automates the targeted illumination process over large numbers of cells or features. At the cellular scale, PSS reveals that oligodendrocyte precursor cells comprise a relatively high proportion of oligodendrocyte-lineage cell population in the cortex compared to the corpus callosum. At the subcellular scale, we selectively sequence the nuclear periphery and demonstrate that the PSS data are consistent with published lamin B1 ChIP-seq data, while also revealing H3K9me3-marked chromatin is enriched at the periphery regardless of laminB1 interaction. Taken together these results demonstrate that PSS has the potential to uncover new connections between spatial and genomic information at diverse length scales.

We anticipate that PSS will be a useful tool for fast, sensitive and robust spatial annotation of cell types and states. In contrast to RNA, DNA is compartmentalized into a cell’s nucleus providing defined regions that are easily identified and well-suited for photoselection. Further, PSS requires the deprotection of two oligonucleotides, which reduces background signal. Finally, large-scale scATAC-seq efforts are underway to create a catalog of accessible regions of the genome, which reflect highly cell-type specific signatures of cell types and states^41^. To demonstrate this, we combine a scATAC-seq atlas with PSS to assess the spatial distribution of cell types within the mouse brain. To do this, we establish a computational strategy which uses topics from scATAC-seq data to deconvolve spatial PSS bulk data. Future work may improve upon this strategy by leveraging recent advancements of bulk sample deconvolution^42^.

The PSS method is broadly accessible to researchers, as all materials are commercially available. PSS is compatible with many imaging systems, including laser-scanning confocal microscopes, targeted illumination devices, and digital micromirror devices. For point scanning devices, spatial resolution of PSS is determined by the point-spread function of the targeted-illumination laser inside the sample volume, which is mainly a function of the numerical aperture of the objective lense and the wavelength of the targeted-illumination laser. On our system, we expect a lateral resolution of approximately 300 nm (Extended Data Fig. 1C), which is sufficiently compact to photoselect sub-cellular regions. We note that the targeted illumination laser passes through the entire thickness of the specimen. Therefore, targeted illumination of small ROIs within the nuclear interior risks background from spuriously unblocked fragments above and below the focal plane. However, this effect may be mitigated by optimizing imaging conditions, for example decreasing the targeted-illumination-laser power such that photocleaving the blocked adapter is only efficient near the center of the light cone. Future iterations of PSS may uncage the libraries using two photon microscopy, which would increase the axial (and lateral) resolution.

The PSS library preparation is compatible with numerous genomic and epigenomic measurements. For example, the *in situ* library preparation stage may be compatible with multiple tagmentation strategies (provided that the transposase carries the photocaged adapters), and we have already documented construction of both ATAC and unbiased whole genome libraries here. An avenue for future exploration would be to generate protein-targeted fragment libraries using a recently-described strategy where proteinA-Tn5 transposase fusion is anchored to the target protein via a specific antibody, resulting in site-specific tagmentation of the underlying chromatin^18,19^. Using photoselection would restrict profiling to specific ROIs, such as particular nuclei, or even subcellular regions. Overall, we expect that the library preparation and downstream analysis phases may be tailored to a wide variety of biological applications. In summary, PSS is a novel spatial genomic method that applies across a diverse range of cell types and species, and is straightforwardly adaptable to accommodate numerous genomic and epigenomic measurements. By combining imaging and sequencing based readouts PSS will uncover new connections at the interface of physical and genomic space. We expect that the user-friendly protocol and lack of specialized materials will facilitate uptake throughout the research community.

## Methods

### Animal Handling

Animal procedures conducted at the Broad Institute complied with the US National Institute of Health Guide for the Care and Use of Laboratory Animals under protocol number 0211-06-18. We housed d (Charles River Laboratory) mice using a 12:12 light-dark cycle with ad libitum access to food and water.

### Mouse Perfusion

Mouse brains were either prepared in-house (cell type benchmarking experiment) or purchased by special request from Zyagen (oligodendrocyte-lineage regional diversity experiment). Purchased mouse brains were obtained from PBS perfused adult mice and shipped flash frozen in accordance with Zyagen’s protocols. For brain tissues prepared at the Broad, C57BL/6 mice were anesthetized in a gas chamber by administration of flowing 3% isoflurane for 1 min. Successful anesthesia was confirmed by a lack of tail-pinch reaction. For the remainder of the procedure, animals were transferred to a dissection tray and maintained under anesthesia using a nose cone flowing 3% isoflurane. To remove blood and protect the tissues during freezing, we perform transcardial perfusion using an ice cold solution containing 10 mM HEPES pH 7.4, 110 mM NaCl, 75 mM sucrose, 7.5 mM MgCl2, and 2.5 mM KCl. Once removed, the brains were frozen for 3 minutes in liquid nitrogen vapor, then transferred to -80ºC for long term storage.

### Cell culture

All cells were cultured at 37ºC with 5% CO_2_ in a humidified incubator. HeLa cells were cultured in high-glucose Dulbecco’s Modified Eagle Medium (DMEM) (ThermoFisher 10569010) supplemented with 10% fetal bovine serum (FBS)(ThermoFisher 16140071) and 1X penicillin-streptomycin (Gibco 15140-122). Similarly, IMR90 human lung fibroblasts were cultured in Eagle’s Minimum Essential Medium (EMEM) (ATCC 30-2003) with the same supplements as above. Cells were maintained in 100 mm sterile TC-treated culture dishes (Falcon 353003) and passaged when they reached approximately 80% confluence. For passaging, cells were washed once with Dulbecco’s phosphate-buffered saline (DPBS) (Sigma D8537) and treated with 0.05% Trypsin-0.02% EDTA (Invitrogen 15575-038) for ∼3-5 minutes at 37ºC. The trypsinized cells were then resuspended in culture media and transferred to a fresh 100 mm plate at a final dilution of ∼1/50.

During passaging, cells for PSS experiments were seeded into glass bottom 96-well plates (Greiner Bio-One 655892) that were pre-treated with Matrigel Matrix (Corning 356234). To prepare the plate, matrigel was diluted 1:50 into ice cold base medium (DMEM/EMEM for HeLa/IMR90) using a refrigerated pipette tip and distributed into the wells (60 µL/well). The plate was incubated for one hour at room temperature before excess solution was aspirated, and the plate was allowed to dry uncovered (in the hood) for an additional 30 min. Cells were seeded for imaging at roughly 55% confluence and placed in the incubator overnight such that they sufficiently recover and adhere before fixation.

### Tissue Sectioning

Tissue samples were placed in a -20ºC cryostat (Leica CM1950) and allowed to equilibrate for 20 min prior to handling. We mounted the tissue onto a cutting block using OCT and proceeded to collect 10 µm thick coronal sections from the desired brain region. For dentate gyrus selection and the oligodendrocyte-lineage heterogeneity experiment, we trimmed away the portion of the brain below the corpus callosum (Extended Data Fig. 2A) using a chilled razor blade. For the oligodendrocyte-lineage benchmarking experiment (Extended Data Fig. 2A), we hemisect the tissue slice by making a sagittal cut along the midline. Tissue sections were manipulated into a 24 well glass bottom imaging plate (Cellvis #P96-1.5H-N or Greiner 655892) and melted onto the surface by briefly placing a finger on the underside of the glass. Plates containing tissue sections were either stored at -80ºC or held briefly in the cryostat before proceeding to fixation. The remainder of the tissue block was returned to the -80ºC for future use.

### General protocol notes

We find that the PSS protocol is streamlined by the use of glass-bottom well plates which support both imaging and enzymatic steps (e.g Greiner 655892). Importantly, we select plates with black well dividers to reduce light scattering and avoid cross-talk during the photoselection stage. We recommend using the smallest possible well size to conserve enzyme during the in situ library preparation, and find that 96- and 24-well formats are suitable for cultured cells and tissue sections, respectively. Unless otherwise specified, we use 100 µL (cells) or 400 µL (tissue) of buffer per well for wash steps, and 50 µL (cells) or 250 µL (tissues) of solution per well for enzymatic steps. Finally, any solutions or samples containing the photocleavable adapters (Extended Data Table 1; PSS1 and PSS2) should be protected from light (e.g. using aluminum foil) whenever possible. All oligonucleotide sequences and antibody information are provided in Extended Data Tables 1 and 2, respectively.

### Fixation and permeabilization (cells)

Cells were rinsed once in 1X PBS to remove excess growth media, then fixed in 1% methanol-free paraformaldehyde (Electron Microscopy Sciences 15714) in 1X PBS for 10 minutes at room temperature. Excess fixative solution was removed by 3×5 min washes in 1X PBS. Cells were permeabilized in 0.5% Triton X-100 (Sigma 93443) in 1X PBS for 10 min at room temperature, then washed 3×5 min in 1X PBS.

### Fixation and permeabilization (tissues)

Tissue sections were covered in 1% methanol-free paraformaldehyde (Electron Microscopy Sciences 15714) in 1X PBS for 10 minutes at room temperature. The fixation reaction is quenched by incubating in 250mM Tris-HCl 8 (Invitrogen 15568) in 1X PBS for 5 min at room temperature. The tissue is washed 3×5 min in 1X PBS. It was then permeabilized with 0.5% Triton-X 100 (Sigma 93443-100) in 1X PBS for 20 min, and washed again three times for five minutes. Fixed and permeabilized tissue was stored at 4ºC in 1X PBS until use.

### Unbiased Tn5 transposase insertion

For unbiased libraries (nuclear periphery experiment), we gently denatured the histones by treating with 0.1 N HCl (Sigma H9892) diluted in water for 5 min at room temperature, then washed 3×5 min in 1X PBS^36^. This step should be completed prior to immunofluorescence, and is not used for ATAC-seq libraries.

### Immunofluorescence

For instances where a fluorescent marker was needed to identify ROIs, we used the following immunofluorescence protocol. Permeabilized cells or tissues were blocked using 4% Bovine Serum Albumin (Sigma A2153) dissolved in 1X PBS with 0.1% Tween 20 (Sigma P9416) (PBST-BSA) for 30 min at room temperature. Samples were stained using a 1:200 dilution of the primary antibody (Extended Data Table 2) in PBST-BSA for either 1 hour at room temperature (cells) or 4ºC overnight (tissues). Excess primary antibody was removed by three five-minute washes in PBSTween (0.1% Tween 20 in 1X PBS) at room temperature. We next incubated the sample in a 1:200 dilution of an appropriate fluorescently-labeled secondary (diluted in PBST-BSA) for one hour at room temperature. Excess secondary was removed by three five-minute washes in 1X PBS. Immunostained samples were stored in 1X PBS at 4ºC for up to a few days before proceeding to the PSS library preparation.

### Purifying Tn5 Transposase

Although Tn5 transposase was purified in-house, it is possible to purchase the enzyme commercially (Lucigen TNP92110) and proceed directly to adapter annealing. Tn5 transposase was purified as previously described with modifications^43^. Briefly, pTBX1-Tn5 (Addgene 60240) was freshly transformed into E. coli competent cells (NEB C3013). A single colony was used to inoculate 10 mL of terrific broth (TB) media with 100 µg/mL carbenicillin, and the culture was incubated at 37°C overnight with shaking. The starter culture was diluted by a factor of ∼100 into fresh pre-warmed TB with 100 µg/mL carbenicillin, then incubated at 37°C with shaking until the OD_600_ reached ∼0.5. Cultures were cooled on ice to room temperature and induced by adding IPTG to a final concentration of 0.3 mM. Once induced, cultures were incubated at room temperature for 16-20 hours with shaking to allow Tn5 transposase expression. Bacteria were harvested by centrifugation at 3700 rpm for 15 min with refrigeration to 4°C. The cell pellet may be stored at -80°C until transposase until purification.

For lysis, 20 g of frozen bacterial cell pellet was resuspended in 200 mL HEGX buffer (20 mM HEPES-KOH pH 7.2, 800 mM NaCl, 1 mM EDTA, 0.2% Triton, 10% glycerol) supplemented with 10 µL of benzonase (Thermo-Fisher Scientific) and one Roche cOmplete protease inhibitor tablet. Cells were mechanically lysed using a microfluidizer (Microfluidics LM20) and the lysate was cleared by centrifugation at 900xg for 30 min at 4°C. To remove E. coli DNA, 5.25 mL of 10% PEI (pH 7) was added dropwise to the lysate while stirring, and the precipitate was subsequently removed by 10 min of additional centrifugation. Tn5 transposase was bound by mixing the cleared lysate with 30 mL of chitin resin (NEB) in a column and rotating end-to-end for 30 min at 4°C. To release the enzyme from the column, 75 mL of HEGX buffer supplemented with 100 mM DTT was added to the column, and the resin was washed by immediately allowing 30 mL of buffer to flow through. The column is sealed and stored at 4°C for 48 hours to release the enzyme from the resin. Eluted Tn5 transposase was concentrated to ∼1.8 mg/mL using centrifuge columns and then dialyzed into 2X storage buffer: 100 mM HEPES-KOH at pH 7.2, 0.2 M NaCl, 0.2 mM EDTA, 2 mM DTT, 0.2% Triton X-100 and 20% glycerol. For long term storage aliquots were held at -80C in 2X storage buffer. The working aliquot was combined 1:1 with 50% glycerol and stored at -20C.

### Anneal Adapters

Photocaged tagmentation adapters (Extended Data Table 1; PSS1 and PSS2) and the blocked mosaic end (Extended Data Table 1; PSS3) were obtained from IDT and resuspended in Ultrapure water (Invitrogen 10977015) to a final concentration of 100 uM (Table 1). The mosaic end sequence was annealed to the adapters using the following reaction: 25 µM PSS1, 25 µM PSS2, 50 µM PSS3, 10 mM Tris-HCl and pH 8.0 and 50 mM NaCl. The annealing reaction was placed in a thermocycler and the temperature was ramped from 85°C to 20°C over approximately 1 hour. Annealed adaptors were mixed 1:1 with glycerol and stored at -20°C until Tn5 loading.

### Load Tn5 Transposase

Annealed adapters were loaded into Tn5 transposase by combining the annealed adapters, Tn5 dilution buffer (50 mM Tris-HCl pH 7.5, 0.1 mM EDTA, 100 mM NaCl, 0.1% NP-40, 1 mM DTT, 50% glycerol) and 16.8 μM Tn5 transposase in a 2:1:1 ratio, then incubating for 30 min at room temperature. Loaded Tn5 was stored at -20°C until use (for up to a few weeks).

### In Situ Library Preparation

Loaded Tn5 transposase was diluted 1:20 (cells) or 1:12 (tissues) in reaction buffer (0.3X PBS, 10 mM Tris pH 7.5, 5% dimethylformamide, and 10 mM MgCl_2_). The solution was applied to the samples and the well plate was sealed with an adhesive film (Applied Biosystems 4306311) to prevent evaporation and protected from light. The plate was incubated at 37°C for 3 hours to allow transposase insertion. The transposition reaction was inactivated by replacing the reaction buffer with 50 mM EDTA in 1X PBS and incubating at 37°C for 30 min. Samples were stained using 5 µM DRAQ5 (Abcam ab108410) in 1X PBS for 5 min and washed once with 1X PBS prior to the photoselection stage.

### Imaging system

Imaging was performed using a Nikon Ti2-E inverted microscope equipped with a Yokogawa CSU-W1 confocal spinning disk unit and a Zyla 2.3 PLUS sCMOS camera. For confocal imaging (to visualize morphological features of the sample and identify ROIs) we used 488 nm, 561 nm and 647 nm laser lines paired with 525/36 (MVI, 77074803), 582/15 (MVI, FF01-582/15-25), and 705/72 (MVI, 77074329) emission filters, respectively. Note that samples harboring PSS libraries should not be imaged using a 405 nm laser line since it will uncage fragments throughout the field of view. For targeted illumination, the microscope was coupled to an XY galvo scanning module via the Ti2-LAPP system (Nikon), and used a 50 mW 405 nm laser line. For targeted illumination, the 405 nm laser power at the objective was set to 1 mW. Samples were imaged through a 1.15 NA CFI Apo LWD Lambda S 40X water immersion objective lens (Nikon MRD77410). The imagining system was controlled by NIS-Elements AR software with the JOBS and General Analysis 3 modules enabled.

### Calibrating the targeted illumination module

Successful targeted illumination requires that (i) the targeted illumination laser is focused in the same plane as the confocal image, and (ii) the position of the steering mirrors is correctly offset such that the beam precisely scans the on-screen ROIs. We note that all commercial targeted illumination devices we tested have mechanisms for adjusting these properties. To ensure these conditions are met, we calibrate the targeted illumination device prior to each photoselection session. To do this, we use a lawn of surface-immobilized fluorophores plated in the same style well-plate as the sample. Specifically, a well is covered with 0.1% poly-l-lysine solution (Sigma P4832) for 10 min at room temperature and then rinsed with Ultrapure water. We bind a fluorescently-labeled PSS adapter (e.g. PSS1 or PSS2) by adding it to the well at 1µM and incubating for 10 min at RT. Rinsing the well twice with Ultrapure water removes unbound oligo. To mitigate any chromatic distortion effects, we advise selecting the same fluorophore for calibration as will be used to outline photoselection ROIs in the sample. We place the calibration plate on the microscope and focus on the lawn of fluorescence in confocal mode. To focus the targeted illumination device, we pulse the laser and inspect the size of the resulting dark spot in the lawn of fluorescence (formed by cleavage and diffusion of the 5’ fluorophore from the PSS adapter). We repeat this process while incrementally adjusting the focus knob on the device until a small and symmetrical spot is achieved. On our system the minimal spot size is roughly 300 nm (Extended Data Fig. 1C). Next, we calibrate the registration between confocal imaging and targeted illumination using the system applet which provides feedback between the mirror position and the laser-spot position in the image. Briefly, this involves steering the beam to five sample points within the confocal image and manually locating the spot (in the lawn of fluorescence) to collect coordinates. Finally, we confirm successful calibration by manually drawing and stimulating ROIs on the lawn of fluorescence and ensuring that the photocleaved regions have crisp edges and are precisely matched to the ROI boundary (Extended Data Fig. 1A).

### Photoselection

The photoselection procedure is summarized as follows: for each relevant field of view within the sample (i) acquire a multichannel confocal image to visualize the fragment library and sample morphology, (ii) manually or algorithmically define ROIs, (iii) selectively illuminate the ROI’s with near-UV light and (iv) acquire a second confocal image to ensure successful cleavage of the PSS adapter, as indicated by loss of fluorescence. We note that it is feasible to implement this process manually for selection of large spatial regions (e.g. the dentate gyrus), but highly recommend a programmatic approach for more numerous regions, especially if they are distributed throughout the sample. Most commercial imaging systems with laser-scanning capabilities are controlled by software equipped with some degree of built-in image processing capabilities. In our hands, NIS Elements AR with the JOBS and General Analysis 3 modules enabled was best suited to PSS, and all photoselection was automated using these tools. The image-processing specifications quoted in the following sections are particular to the adjustable parameters in our Nikon software, however the process is generalizable to other systems.

For the HeLa cell demonstration (Figure 1B), we identified nuclei using the fluorescence signal from the inserted PSS adapters (PSS1 and PSS2) which carry the Alexa 546 fluorophore. Specifically, we acquired a single-plane confocal image (in the plane with the highest contrast) using a 561 nm laser line paired with a 582/15 emission filter. A binary mask of the nuclei was generated by applying a low pass filter (5X), a local contrast filter (size 60, power 40%), and then thresholding the pre-processed image, with the “fill holes” and “separate objects” options selected. An appropriate threshold level was determined by visually comparing the outline of the binary mask to the raw confocal image. We manually selected a subset of regions from the binary mask (nuclei) to targeted illumination ROIs. To generate the targeted illumination movie (Supplemental Movie 1, Fig 1B shows first and last frames), we performed confocal imaging (651 nm laser, 582/15 filter), while simultaneously scanning the 405 nm laser through the ROIs (300 µsec dwell time).

For granule cell selection (Fig 2B, top), we roughly located the dentate gyrus region of mouse hippocampal brain slices stained with DRAQ5 by scanning the entire sample using a 10X objective (Nikon MRD00105) with 640 nm illumination and a 705/72 emission filter. We switched to the 40X objective and generated a tiled grid of (∼20) XY positions (1% overlap) over the entire dentate gyrus region. For each XY position, we automatically identified nuclei belonging to the dentate gyrus by acquiring a single-plane confocal image (DRAQ5 stain, 640 nm laser, 705/72 emission filter). We created an initial binary mask of all nuclei applying a low pass filter (5X) followed by thresholding. We used the comparatively high density of nuclei in the dentate gyrus region compared to the surroundings to distinguish the target regions. Specifically, we dilated the initial binary mask using a 10 µm radius disk such that the individual nuclear masks in cell-dense regions merged into a large single region while nuclear masks from sparse regions did not merge. We eroded the binary using a 10 µm disk and filtered out small regions by gating on area. To generate the final masks for targeted illumination we performed an and operation on the initial and processed masks. We note that in this instance, hand drawing the boundaries is adequate and produces similar results.

We identify oligodendrocyte-lineage cells (Fig 2B bottom) using a DRAQ5 DNA stain paired with an immunostain against OLIG2 (Alexa 488 secondary). We generated a tiled grid (1% overlap) spanning the entire brain slice and looped over the (∼200) XY positions using JOBS. At each XY position we performed the following steps:

- Run an autofocus using the DRAQ channel to select the plane with the highest contrast
- Capture confocal images of the nuclei (DRAQ5), OLIG2 immunostain (Alexa 488), and ATAC library (Alexa 546)
- Generated nuclear masks on the DRAQ5 image by applying a low pass filter (5x), local contrast filter (size 60, power 40%) and thresholding with with the “fill holes” and “separate objects” options selected
- Refine the mask to include only oligodendrocyte-lineage nuclei by gating on the mean fluorescence intensity of the OLIG2 immunostain
- Perform targeted illumination using the boundaries defined by the mask above
- Capture a confocal image of the ATAC library to ensure successful cleavage in oligodendrocyte nuclei

For the oligodendrocyte-lineage heterogeneity experiment, we identified the corpus callosum and cortex regions (in separate brain slices) visually from a 10X scan of the sample (e.g. Extended Data Fig. 2D). We manually drew masks for these regions before switching to 40X for the targeted illumination phase. We performed the same procedure described above, except that we took the logical ‘and’ of the region mask and the oligodendrocyte-lineage nuclei mask prior to photostimulation to restrict sequencing data collection to the desired region.

For the nuclear periphery experiment (Figure 3), IMR90 cells imaged using a DRAQ5 DNA stain and an immunostain against LamB1 with an Alexa 488 secondary. We generated a 17×17 grid of XY positions with 1% overlap, which covered the majority of the well. We looped through the XY positions using the JOBS module, and performed the following steps:

- Run an autofocus using the DRAQ channel to select the plane with the highest contrast
- Capture confocal images of the nuclei (DRAQ5), LamB immunostain (Alexa 488), and ATAC library (Alexa 546)
- Generate nuclear masks on the DRAQ5 image by applying a low pass filter (5x), local contrast filter (size 60, power 40%) and thresholding with with the “fill holes” and “separate objects” options selected
- Remove regions touching borders
- Filter out cells with convexity < xx and circularity <yy to such that only healthy cells are included in analysis
- Erode the nuclear mask using a 300 µm radius disk to generate an inner mask
- Subtract the inner mask from the nuclear mask to generate ring-shaped ROIs at the nuclear periphery
- Perform targeted illumination using inside the regions bounded by the mask above
- Capture a confocal image of the ATAC library to ensure successful cleavage

### Reverse crosslinking and library purification

Samples were digested by incubation in reverse-crosslinking buffer (50 mM Tris pH 8, 50 mM NaCl, 0.2% SDS) with 1:50 proteinase K (NEB P8107S) for 8-16 hours at 55°C. The digestion buffer was added directly to the well plate, which was sealed with an adhesive film (Applied Biosystems 4306311) and protected from light. Digested samples were column purified using the NucleoSpin Gel and PCR Clean-Up kit (TaKaRa 740609) according to the manufacturer’s directions, except we perform two successive rounds of elution using 20 µL then 16 µL of 10 mM Tris pH 8 (regardless of well size). At this stage, we optionally reserve half the eluate volume to create an input library (i.e. amplify all fragments regardless of if they were previously unblocked) to enable enrichment-based measurements in the analysis stage. Specifically, the eluate from each sample is split into two PCR tubes, each containing 16 uL of volume. We place one one of these PCR tubes under the collimated light from a 365 nm LED (LED: Thorlabs M365LP1-C5, driver: Thorlabs LEDD1B) for 3 minutes such that all fragments become uncaged. The power per area inside the LED spot is 0.15 mW/mm^2^. Input libraries are then processed in the same manner as the photoselected libraries in all subsequent steps.

### Library structure

PSS libraries were designed to share essential features with Illumina Nextera libraries and eliminate the need for custom primers during the sequencing phase. We note that the PSS tagmentation adapters (ISS1 and ISS2) are identical to the Nextera read 1 and read 2 adapters except that they are modified on the 5’ end with a photocleavable spacer and fluorophore. The central principle of PSS is that complete sequenceable fragments are only generated if the tagmentation adapters are photocleaved, exposing a 5’ phosphate group. To this end, we ligate secondary adapters that carry the Illumina P5 and P7 sequences to the ends of uncaged fragments. To ensure correct pairing with the tagmentation adapters (P5 with adapter 1 and P7 adapter 2), secondary adapters are assembled with the fragment library using splint oligos to facilitate the ligation reaction. For demultiplexing after sequencing, the P7 secondary adapter additionally carries an 8 bp index sequence followed by an added constant region for splint binding (Extended Data Fig. 1A). Libraries are ultimately amplified for sequencing using primers that bind the P5 and P7 sequences on the secondary adapters. A fully detailed version of the library structure is shown in Extended Data Fig. 1A.

### Adapter ligation

We ligate secondary adapters (PSS4 and PSS5) to the ends of uncaged fragments using splints (PSS6 and PSS7) to stabilize the assembly (Figure 1A, step 4; Extended Data Fig. 1A; Extended Data Table 3). We first perform an annealing step to transiently pre-assemble fragments by adding 4 µL of annealing master mix (2.5 µL of 10X T4 Ligase Buffer, 0.25 µL of ISS4, 0.25 µL of 100 µM ISS5, 0.25 µL of 100 µM ISS6, and 0.75 µL of water) to the 16 µL eluate volume from each sample. Finally, we add 2.5 µL of uniquely indexed ISS5 to each sample (total volume 22.5 µL) and perform the annealing by ramping the temperature from 50°C to 43°C for 10 min, then down to 25C, using a ramp rate of -0.1C/s. Once the temperature reaches 25C, we spike in 2.5 µL of 3M U/mL T7 DNA ligase (NEB M0318L) and continue incubating for 1 hour at 25C, followed by a 10 min inactivation step at 65°C and a 4°C hold. The ligation reaction is purified by adding 25 µL of Ampure XP beads (Beckman Coulter A63880) to the reaction (1:1 ratio) and continuing the clean-up according to the manufacturer’s directions. For elution, we add 23 µL of 10 mM Tris pH8 to the beads such that we reliably recover and transfer 20 µL to a fresh PCR tube.

### Library amplification

The library amplification stage proceeds roughly as described in Buenrostro et al, 2015^17^. In summary, the 20 µL of eluate from the previous step was combined with 2.5 µL of 25 µM PSS8, 2.5 uL of 25 µM PSS9 and 25 µL of NEBNext High-Fidelity 2X PCR Master Mix (NEBM0541L). We cycle the PCR reaction using the following conditions: 72°C for 5 min, 98°C for 30s, 5 cycles of 98°C for 10s then 72°C for 1 min 30s, 72°C for 2 min, 4°C hold. Note that the initial extension step is critical to producing complete double-stranded fragments that can be amplified by PCR (Fig 1A, step 4). To minimize the effects of PCR bias, we determine an appropriate number of additional amplification cycles using qPCR. For a single qPCR reaction, we combine 5 uL of previously amplified DNA with 0.25 µL of 25 uM PSS8, 0.25 µL of 25 µM PSS9, 0.09 µL of 100X SYBR Green, 5 µL NEBNext High-Fidelity 2× PCR Master Mix, and 4.41 uL of water. We typically run two replicate reactions for each sample, and store the remaining 40 µL of partially amplified DNA at 4°C during the qPCR. The reactions are cycled in a qPCR instrument using the following conditions: 98°C for 30 sec, 20 cycles of 98°C for 10 sec then 72°C for 1 min 30 s, 72°C for 1 min. The calculated number of additional cycles corresponds to the point where the relative fluorescence reaches 25%-30% of the maximum value.

At this stage, it can be useful to visualize the qPCR reaction on an agarose gel. This serves as an initial quality control and can detect the lengths of any dimers that should be removed by size selection during the eventual PCR clean up. To do this, we fill the sample wells of a 2% E-Gel EX Agarose Gels (ThermoFisher G401002) with 16 µL of Ultrapure water and 4 µL of qPCR reaction. The marker well is filled with 20 µL of a 1:40 dilution of 100 bp ladder (NEB N3231L). The gel is run on an E-Gel Power Snap Electrophoresis Device (ThermoFisher G8100) according to the manufacturer’s directions. Once the run is complete, we verify that a library is present and inspect the lanes for dimer. We typically observe a few bright bands that are shorter than or equal to the 100 bp marker, and use this information to guide our choice of a 1.2x ratio of Ampure XP beads to sample volume during the clean-up stage. An example gel is shown in Supplemental Figure XX.

The original partially amplified libraries are returned to the thermocycler for the calculated number of cycles (N) using the following program: 98°C for 30s, N cycles of 98°C for 10 sec then 72°C for 1 min 30 sec, 72°C for 2 min, 4°C hold. PCR reactions are cleaned by adding 48 µL of Ampure XP beads (Beckman Coulter A63880) to the 40 µL reaction volume (1.2x ratio), then following the manufacturer’s directions. Libraries are eluted in 20 µL of 10 mM Tris pH8 and stored at -20°C until quantification and sequencing.

### Library Quantification

Libraries are quantified via qPCR using the KAPA Library Quantification Kit (Roche 0796020400), which specifically analyzes sequencing-competent fragments. We run 1:10000 and 1:1000000 dilutions of the original library and follow the manufacturer’s procedure for quantification and analysis. The mean fragment length is determined using a Bioanalyzer (Alegent) following the manufacturer’s directions, however we do not find it necessary to repeat this process during every library preparation. In our hands, using a mean fragment length of 750 bp to calculate the molarity of the library consistently produces an appropriate clustering density during sequencing.

### Sequencing

Libraries were sequenced on an Illumina Nextseq using the following read structure: 60 bp read 1, 60 bp read 2, 20 bp index 1.

### Sequencing alignment and pre-processing

Sequenced fragments were trimmed for adapter reads and aligned to either the mm10 (mouse brain experiments) or hg38 (HeLa or IMR90 experiments) reference genomes using Bowtie2^44^. Bowtie was called with the default parameters, except ‘-X 2000’ was set to allow alignment of fragments up to 2000 bp. Aligned fragments were sorted by genomic position and filtered using alignment quality (MAPQ > 30 retained). Alignments to unlocalized and unplaced sequences, the Y chromosome, and the mitochondrial DNA were excluded from downstream analysis. Duplicate reads were removed using the Picard tools MarkDuplicates function (http://broadinstitute.github.io/picard), which also provides an estimate for the calculated library size using the duplication rate.

TSS enrichment scores (e.g. Fig. 1C) were calculated for ATAC-seq libraries as previously described in Buenrostro et al. 2013^20^. In short, we obtained bed files containing mm10 and hg38 TSS locations using the UCSC table browser. Fragments aligned within 2000 bp of a TSS were included in analysis, and the enrichment score was taken as the ratio of the number of fragments at each relative position and the mean number of fragments observed in the regions 1800-2000 bp of TSSs genome wide.

### Signal to noise calculation

To estimate the signal-to-noise ratio achieved by the PSS blocking mechanism (Fig. 1D), libraries were constructed either in HeLa cells or tissue slices. Each sample was digested immediately and the eluate was split in half, generating a pair of blocked libraries with equal numbers of fragments. One library from each pair was unblocked under a 365-nm LED (LED: Thorlabs M365LP1-C5, driver: Thorlabs LEDD1B) for 3 minutes (cleaved half), while the other library was protected from light (blocked half). All libraries were processed as described above until an estimation of the library size (calculated number of unique fragments) was reached (see sequencing alignment and pre-processing). The percent signal is calculated as follows:

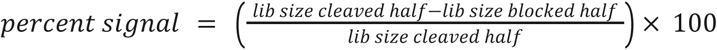

We note that the small number of fragments detected in the blocked libraries likely originate from (i) incompletely synthesized oligos that are contaminated with exposed phosphate groups, (ii) a low rate of unintentional cleavage in the room light, and (iii) misassignment of sequencing reads during the demultiplexing stage.

### Smoothed coverage traces

Smoothed coverage traces (Fig. 1C and Fig. 2B) were generated as follows. For each paired-end alignment, reads on the forward strand were shifted by +4 bp and reads on the reverse strand were shifted by -5 bp to account for the 9 bp duplications generated by Tn5 transposase insertions^20,45^. To smooth the appearance of the final coverage traces, fragments were extended by 150 bp on each end. Finally, the adjusted fragment positions were output as a bed file and the bedtools genomecov function was used to generate coverage traces for visualization^46^.

### cisTopic Analysis

Cis-regulatory topics representing 200 distinct axes of biological variation were calculated on the counts matrix of previously published single-cell ATAC-seq data in the mouse brain using cisTopic^23,24^, binarized, and then used as custom annotations within chromVAR^47^ to score oligodendrocyte PSS ATAC-seq counts using the same peak set. Correlation heatmaps were generated by calculating the correlation of z scores of each cisTopic between bulk replicates and pseudobulk clusters from the single-cell data using the pheatmap package.

### Stepwise Linear Regression

Linear regression was performed using the lm function of the stats package in R with cisTopic z-scores of the pesudobulk scATAC-seq profiles of the annotated brain clusters used as independent variables for each bulk PSS ATAC-seq replicate. Variables for the best-performing model for each sample were selected through forward selection using the step function of the stats package in R. Non-negative coefficients were normalized to add up to 1, and then replicates for each condition were averaged to come up with the final cell-type composition estimates.

### LamB ChIP-seq

LamB1 ChIP-seq data were originally published in Dou et al. 2015 and obtained from the NCBI Gene Expression Omnibus (GEO) database under accession number GSE63440^35^. Two replicate LamB1-ChIP-seq and paired input libraries were downloaded and processed as described above (sequencing alignment and pre-processing), except with minor modifications to accommodate single-end reads. Specifically, Bowtie2 was called with the default parameters, and in the filtering stage we removed the requirement that reads map in a proper pair.

### Enrichment traces

For the nuclear periphery profiling experiment (Fig 3), each PSS and ChIP-seq replicate is associated with paired targeted and input libraries (processed as described above). To generate the chromosome level enrichment plot (Fig 3B), the number of reads mapping to each assembled chromosome was queried by running the ‘samtools idxstats’ command on the filtered bam filles (see sequencing alignment and pre-processing). The read depth for each library was taken as the sum of mapped reads across all assembled chromosomes. We normalized the reads per chromosome in the targeted and input libraries by read depth and reported the ratio of these quantities as the enrichment.

To generate the enrichment traces (Fig 3C), the genome (hg38) was divided into 100 kb bins. The targeted and input mapped sequencing reads from all replicates were pooled into single bam files. Raw read counts were generated for the targeted and input libraries using the chromVar ‘getcounts’ function to read the filtered bam files into the genomic bins^47^. Raw read counts were normalized by read depth on a per chromosome basis, where the chromosomal read counts were obtained using ‘samtools idxstats’. An enrichment value for each bin is obtained by taking the ratio of the normalized read counts in the targeted and total libraries. The log (base 2) of the enrichment is displayed.

### Correlation analysis

To generate the relative p-q arm position track for correlation analysis (Fig 3D), we begin with the same 100 kb genomic bins as the enrichment traces. Centromere regions were first obtained from the UCSC table browser, and the midpoint of each region was called as the centromere position. For all chromosomes, a relative p-q arm position was assigned to each 100 kb genomic bin by subtracting the centromere position from the bin center position and normalizing the result by the chromosome length.

All available histone modification ChIP-seq data (and associated inputs) collected in IMR90 cells were downloaded from the ENCODE database (encodeproject.org)^40^. Data were obtained as aligned reads (hg38 bam files) and all post-alignment filtering steps were applied (see Sequencing alignment and preprocessing). Binned enrichment traces ENCODE data were calculated as described above (see Enrichment traces). The H4k16ac ChIP-seq data were obtained separately from Rai et al.,2014^48^, and were processed according to our methods from alignment to enrichment trace generation.

The PSS enrichment trace was correlated with the LamB1 ChIP-seq trace, the relative p-q arm position trace and all of the histone modification enrichment traces on a per-chromosome basis. Traces were retained for further consideration if (i) the magnitude of the chromosome-averaged Pearson correlation coefficient was at least 0.4 and (ii) the magnitude of the Pearson correlation coefficient for at least one chromosome was greater than 0.6. All positively correlated traces that meet these conditions, as well as the 7 most negatively correlated traces are ultimately displayed in the figure. All p-values associated with correlation coefficients shown in the figure are significant. The PSS residuals were calculated using stepwise linear regression (Matlab stepwiselm function), and then correlated with the histone modification ChIP-seq profiles as described above.

### H3K9me3 radial intensity profile

IMR90 cells were immunostained for H3K9me3 (Abcam ab8898) using an Alexa 488 secondary (see Immunofluorescence), and DRAQ5 stained to visualize the nuclei. Single-plane images of both stains were acquired using a 40X objective (see imaging). To calculate the radial intensity profile, nuclear masks were obtained by smoothing the DRAQ5 image using a 3-pixel radius Gaussian blur and thresholding the smoothed image to generate a binary mask. Sub-nuclear-sized objects were filtered from the binary as needed. Next, the H3K9me3 image was background corrected by subtracting the median pixel intensity outside the nuclear masks from each image pixel. The radial intensity (from the H3K9me3 immunostain) was quantified by successively eroding the nuclear mask using a 3-pixel radius disk. At each subsequent erosion step, the inner mask was subtracted from the outer mask, generating a series of hollow regions of decreasing radius (Extended Data Fig. 4C). The median pixel intensity inside each hollow region was quantified and plotted against the normalized radial position, where the positions of the innermost and outermost regions were set to 0 and 1, respectively. To calculate the mean radial intensity, we binned the relative radial positions (0.05 bin size) and averaged the intensity values within each bin.

## Acknowledgments

We are grateful to Jonathan Strecker for assistance with Tn5 transposase Purification. We thank Zack Chiang for providing scripts for the generation of smoothed coverage traces, and for advice on computational methods. F.C. acknowledges support from NIH Early Independence Award (1DP5OD024583), the NHGRI (R01HG010647), the Burroughs Wellcome Fund CASI award, and the Merkin Institute.

## Data availability statement

All PSS data generated in this study will be submitted to an appropriate repository upon manuscript acceptance. Single-cell ATAC-seq data (mouse hippocampus) from Sinnamon et al.^22^ are available through Gene Expression Omnibus (GEO; https://www.ncbi.nlm.nih.gov/geo/) under the accession GSE118987. Single-cell ATAC-seq data (whole mouse brain) from Lareau et al.^23^ were deposited at GEO under accession number GSE123581. All Histone ChIP-seq data are available through the ENCODE encyclopedia^40^ (https://www.encodeproject.org/) except for the H4K16ac data^48^, which are available under GEO accession number GSE56307. Lamin B1 ChIP-seq data from Dou et al.^35^ are available under GEO accession number GSE63440.

## Code availability statement

Analysis scripts will be made available upon manuscript acceptance.

## Extended Data

**Extended Data Table 1.**
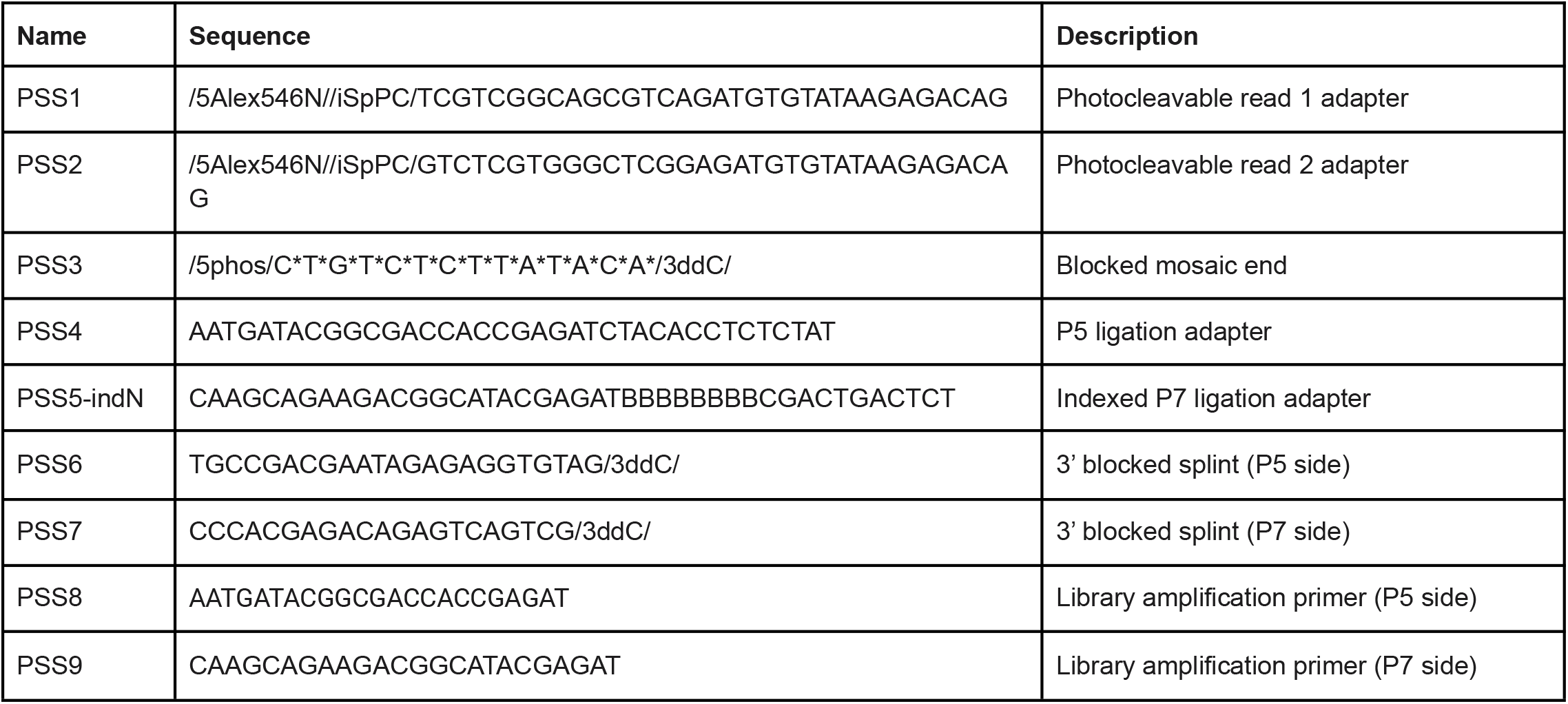
List of oligos used in this study written 5’ to 3’ using IDT modification codes.The B bases in the PSS5-indN primer represent a known barcode sequence for sample demultiplexing. For convenience, our complete set of indexed P7 ligation adapters is provided in Extended Data Table 3.

**Extended Data Table 2.**
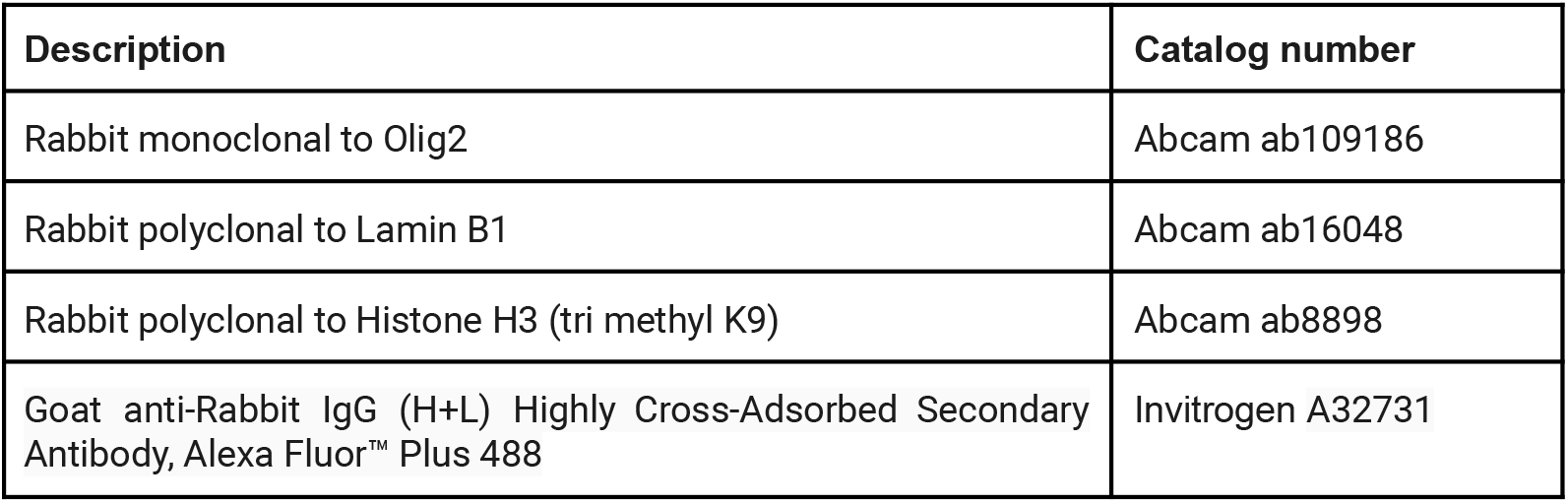
List of antibodies and catalog numbers

| Description | Catalog number |
| --- | --- |
| Rabbit monoclonal to Olig2 | Abcam ab109186 |
| Rabbit polyclonal to Lamin B1 | Abcam ab16048 |
| Rabbit polyclonal to Histone H3 (tri methyl K9) | Abcam ab8898 |
| Goat anti-Rabbit IgG (H+L) Highly Cross-Adsorbed Secondary Antibody, Alexa Fluor™ Plus 488 | Invitrogen A32731 |

Extended Data Table 3- complete list of indexed P7 ligation adapters. Barcode sequences are at least 3 hamming distances apart.

Supplemental Movie 1 - movie demonstrating the photoselection process for ATAC-seq libraries HeLa cells (related to Fig. 1B). Magenta color represents the fluorescence intensity of the photocleavable PSS adapter. Nuclei outlined in blue were exposed to near-UV light using targeted illumination.

**Extended Data Figure 1.**
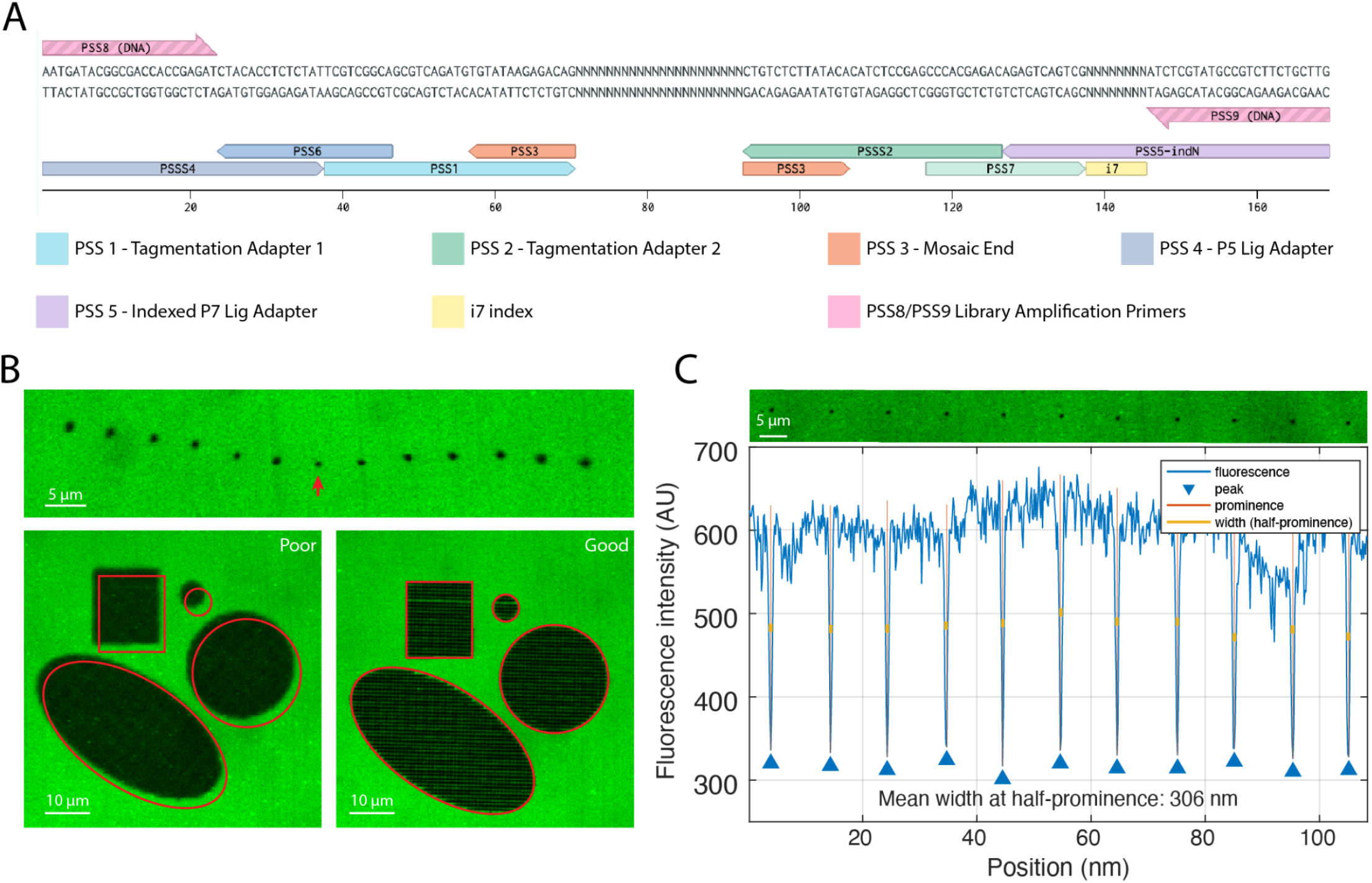
Targeted illumination calibration and detailed library structure. A) Detailed PSS library structure. Central N bases represent the genomic DNA fragment. B) The focus and registration of most targeted illumination devices are tunable. Top image strip shows the photocleaved area (lawn of 488 fluorescence) resulting from a single stimulation point as the instrument is moved through a range of focal positions. Red arrow indicates correct focus. Lower left image shows poor focus (fuzzy edges) and poor registration (shift) while the lower right image shows proper calibration. C) Measuring the maximum resolution of PSS. Image strip shows a line of individual stimulation points on a photocleavable lawn of fluorescence. Spots were measured using the same 405 nm laser power as for PSS experiments (∼1 mW at objective). The fluorescence intensity (blue line) was measured by taking the minimum pixel value of the columns in the image strip above. The minimal spot size is taken as the full width at half-prominence (yellow bars).

**Extended Data Figure 2.**
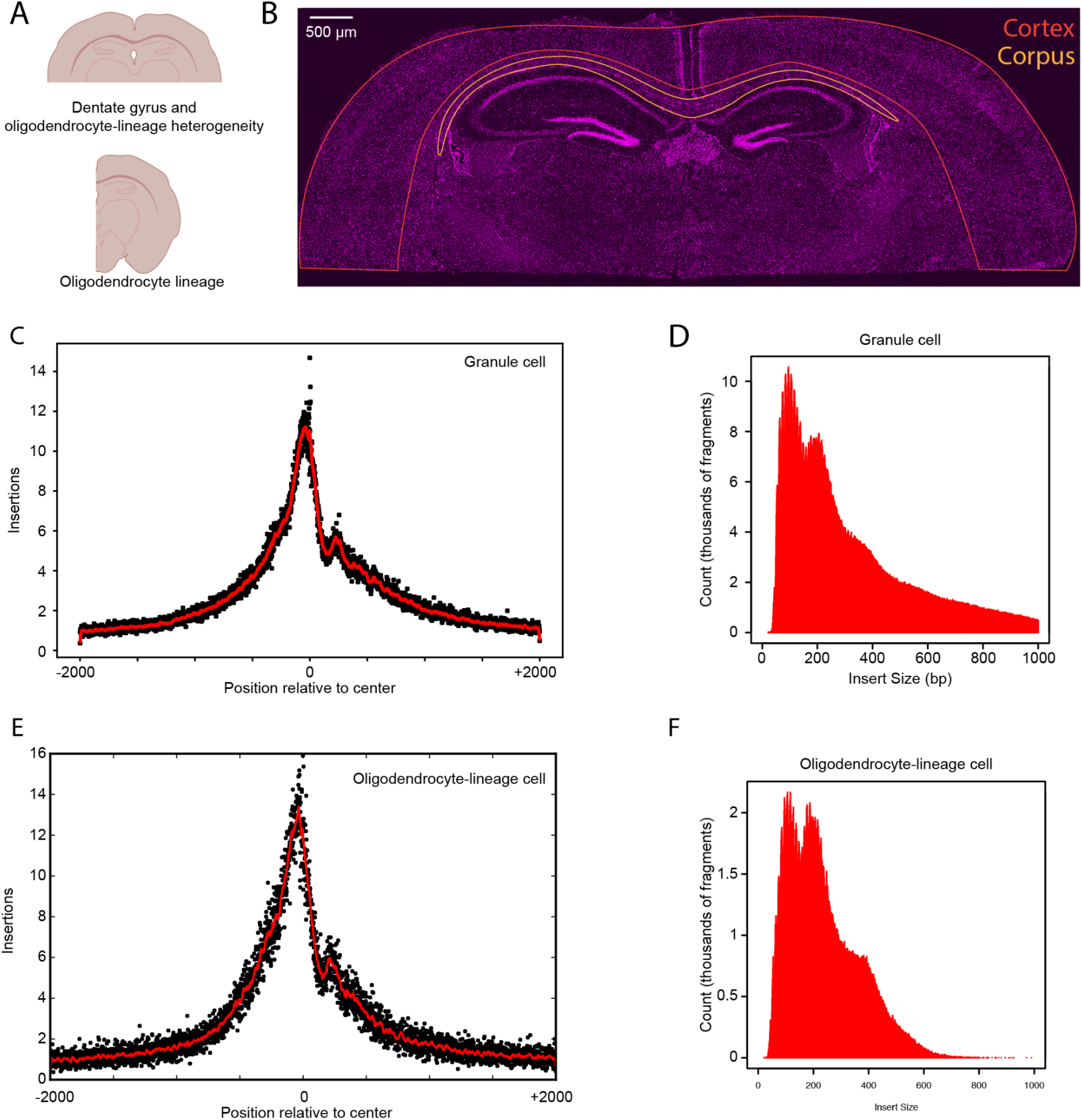
PSS in the mouse brain. A) Schematic diagram shows how hippocampal brain slices were trimmed by experiment. For the dentate gyrus region selection, and the region-specific oligodendrocyte experiments the coronal section was trimmed below the corpus callosum (top). For general oligodendrocyte-lineage selection, the section was subdivided vertically at the midline. B) Low magnification scan of mouse hippocampal region with cortex (red) and corpus callosum (orange) regions outlined. Magenta color represents the fluorescence intensity from DRAQ5 DNA stain. C) Example TSS enrichment plot for PSS granule cells for a single replicate. D) Insert size distribution from a single replicate of PSS granule cells. E) Example TSS enrichment plot for single replicate of PSS oligodendrocyte-lineage cells. F) Example insert size distribution for single replicate of PSS oligodendrocyte-lineage cells.

**Extended Data Figure 3.**
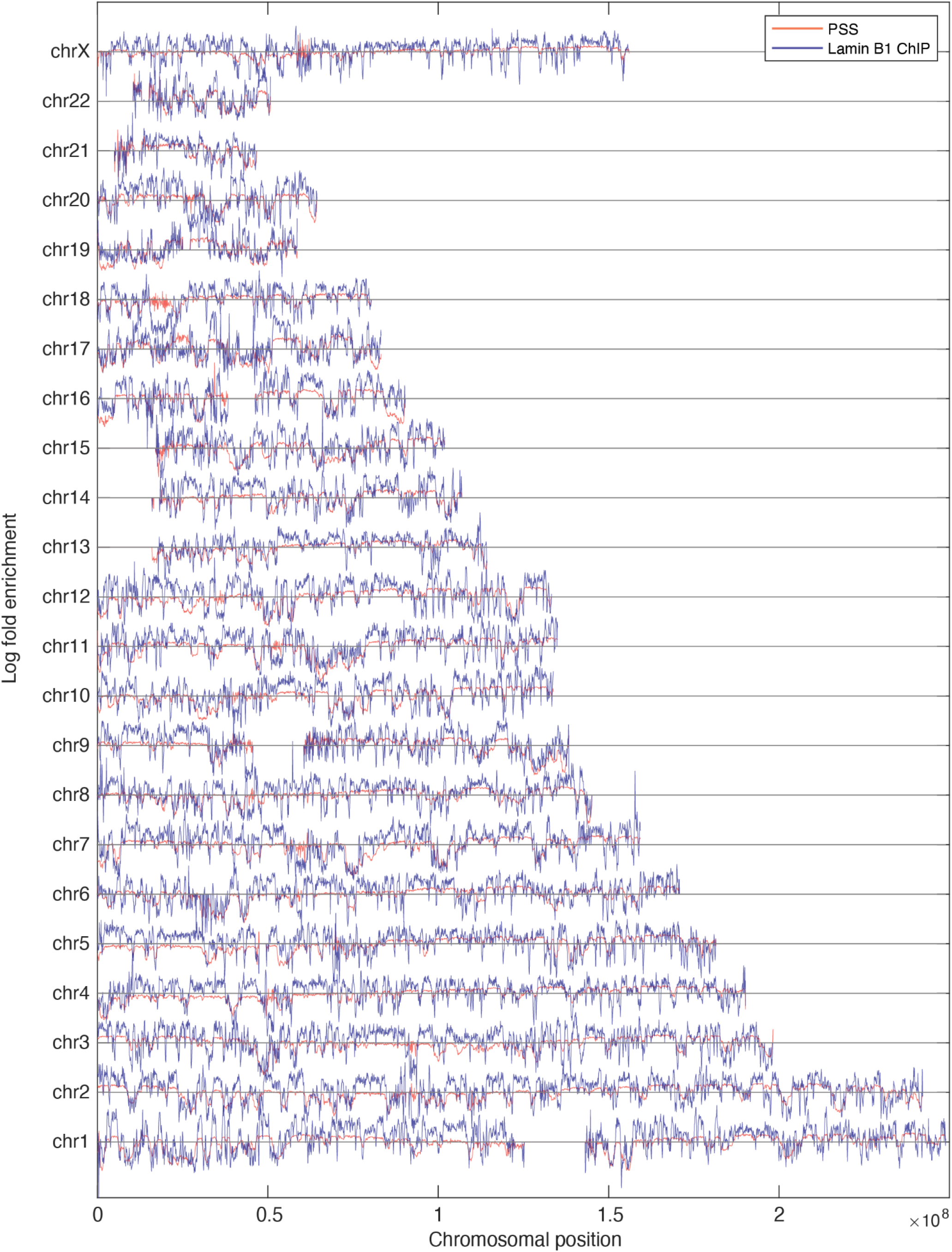
Comparison of PSS and Lamin B1 ChIP-seq profiles for all chromosomes. Orange and blue lines show PSS and ChIP log2 enrichment profiles, respectively.

**Extended Data Figure 4.**
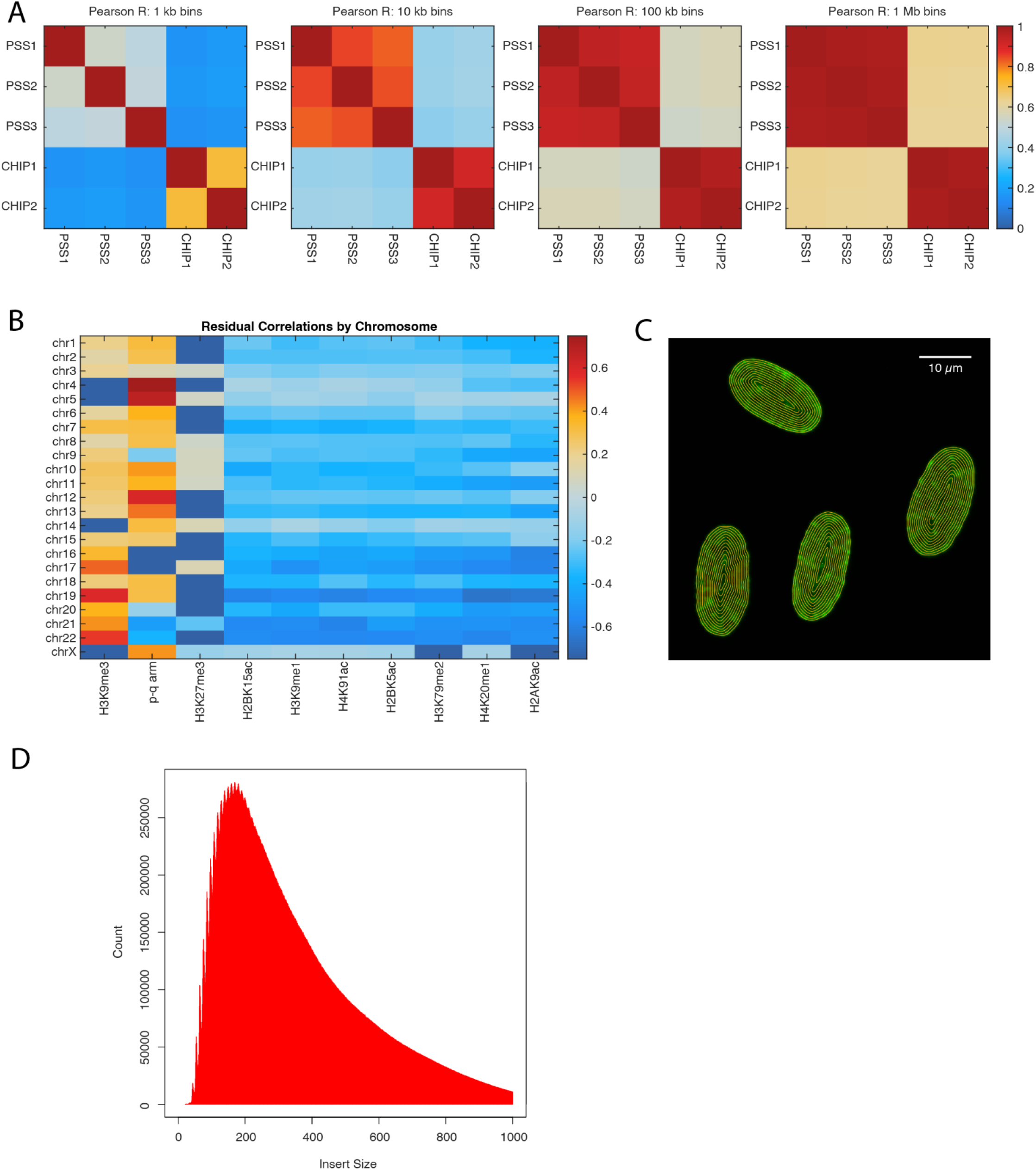
PSS at the nuclear periphery. A) Heatmaps showing pairwise Pearson correlation coefficients for all individual PSS and lamin B1 ChIP-seq replicates at various bin sizes. B) Correlation between PSS residuals (lamin B1 ChIP signal removed) and histone marks from the ENCODE database. C) Images showing ring-like regions for calculating H3K4me3 immunofluorescence intensity as a function of radial position. D) Size distribution of sequenced fragments PSS at the nuclear periphery. Distribution shows a single peak, indicating a loss of sensitivity to histone positioning.

